# Vascularized Brain Assembloids with Enhanced Cellular Complexity Provide Insights into The Cellular Deficits of Tauopathy

**DOI:** 10.1101/2023.06.30.547293

**Authors:** Simeon Kofman, Xiaohuan Sun, Victor C. Ogbolu, Larisa Ibric, Liang Qiang

**Author notes:** Send correspondence to: Liang Qiang, MD/PhD, Department of Neurobiology and Anatomy, Drexel University College of Medicine, 2900 Queen Lane, Philadelphia, PA 19129. co-first author (Simeon Kofman, Xiaohuan Sun).

## Abstract

Advanced technologies have enabled the engineering of self-organized 3-dimensional (3D) cellular structures from human induced pluripotent stem cells (hiPSCs), namely organoids, which recapitulate some key features of tissue development and functions of the human central nervous system (CNS). While hiPSC-derived 3D CNS organoids hold promise in providing a human-specific platform for studying CNS development and diseases, most of them do not incorporate the full range of implicated cell types, including vascular cell components and microglia, limiting their ability to accurately recreate the CNS environment and their utility in the study of certain aspects of the disease. Here we’ve developed a novel approach, called vascularized brain assembloids, for constructing hiPSC-derived 3D CNS structures with a higher level of cellular complexity. This is achieved by integrating forebrain organoids with common myeloid progenitors and phenotypically stabilized human umbilical vein endothelial cells (VeraVecs™), which can be cultured and expanded in serum-free conditions. Compared with organoids, these assembloids exhibited enhanced neuroepithelial proliferation, advanced astrocytic maturation, and increased synapse numbers. Strikingly, the assembloids derived from hiPSCs harboring the tau^P301S^ mutation exhibited increased levels of total tau and phosphorylated tau, along with a higher proportion of rod-like microglia-like cells and enhanced astrocytic activation, when compared to the assembloids derived from isogenic hiPSCs. Additionally, they showed an altered profile of neuroinflammatory cytokines. This innovative assembloid technology serves as a compelling proof-of-concept model, opening new avenues for unraveling the intricate complexities of the human brain and accelerating progress in the development of effective treatments for neurological disorders.

**Significance Statement:** Modeling neurodegeneration in human *in vitro* systems has proved challenging and requires innovative tissue engineering techniques to create systems that can accurately capture the physiological features of the CNS to enable the study of disease processes. The authors develop a novel assembloid model which integrates neuroectodermal cells with endothelial cells and microglia, two critical cell types that are commonly missing from traditional organoid models. They then apply this model to investigate early manifestations of pathology in the context of tauopathy and uncover early astrocyte and microglia reactivity as a result of the tau^P301S^ mutation.

## Introduction

The advent of human brain organoids as relevant *in vitro* and *ex vivo* models has revolutionized neuroscientific research, offering neuroscientists a remarkable opportunity to investigate neurodegeneration alongside animal models. Human induced pluripotent stem cells (hiPSCs) that can be developed into self-organized 3D brain organoids have been used to study cellular and molecular changes stemming from various neurodegenerative diseases.^1^ These advanced and versatile organoid models have undergone remarkable advancements, encompassing diverse cellular profiles and physiological environments, enabling the utilization of innovative technologies to explore disease mechanisms and uncover new insights at unprecedented levels of cellular and molecular resolution. While neurons remain the most prolific cell type and target for study in the field, many non-neuronal cell types of the central nervous system (CNS), including astrocytes, microglia, and vascular endothelial cells, have emerged as essential mediators of pathology and warrant devoted attention in the study of these diseases.^2^ Despite some recent innovations, the majority of existing organoid technologies still exhibit a critical limitation by lacking essential CNS components, specifically microglia and endothelial cells. Some successful attempts have been made to integrate these cell types, but the availability of models incorporating microglia and endothelial cells remains limited, and their application in modeling neurodegenerative diseases is yet to be fully explored.^1^ This study presents a groundbreaking CNS assembloid model that integrates hiPSC-derived forebrain neurons and astrocytes together with hiPSC-derived microglia-like cells and VeraVecs™, a human umbilical vein endothelial cell (HUVEC) with enhanced culture durability, to offer a comprehensive representation of the complex cellular milieu. Furthermore, we decide to assess the efficacy of this innovative system within the context of primary tauopathies, where both the immune and vascular components of the CNS play crucial roles. With an unwavering focus on tauopathies, we aim to unravel the neuropathological intricacies inherent to these debilitating conditions, while assessing the fidelity and translational potential of our innovative model in faithfully replicating some of the essential pathological alterations.

## Methods and Materials

The detailed materials and methods are depicted in the supplementary information.

## Results

The hiPSCs used in the generation of organoids and assembloids were validated for pluripotency with markers TRA-1-60 and TRA-1-81, as well as their potential of three germ layer differentiation with SOX17, Brachyury, and OTX2. (Figures 1A-I). hiPSCs were aggregated into embryoid bodies (EBs) and cultured in neuronal induction media supplemented with dual SMAD inhibition, followed by the optimized protocol (Figure 1J). These organoids were characterized for populations of neural progenitor cells (NPCs), neurons, and astrocytes (Supplements 1A-D). Note that the presence of oligodendrocytes was limited in number in these organoids (data not shown). Vascularized assembloids were generating by dissociating 3-week-old forebrain organoids and re-aggregating with hiPSC-derived common myeloid progenitors (CMPs) and VeraVecs™ in an AggreWell plate (Figure 1K). We also attempted to generate assembloids by dissociating organoids cultured for less than 3 weeks or more than 3 weeks; however, the former resulted in insufficient neuroectodermal induction with inadequate neuronal and astrocytic populations, while the latter led to significant cell death during the re-aggregation process (data not shown). We used VeraVecs™, which are HUVECs engineered with transgenic exon4 open reading frame 1 of the adenovirus (E4ORF1), to introduce a durable endothelial cell component into the model. The E4ORF1, an adenoviral gene product that maintains endothelial cells in an early-passage state, permits both phenotypic stability and improved survival in serum free conditions.^3^ E4ORF1, in combination with vascular endothelial growth factor (VEGF), is also critical for maintaining ECs’ angiogenic potential.^3^ Before integrating into assembloids, red fluorescent protein (RFP)-labelled VeraVecs™ were shown to express endothelial cell marker CD31 and tight junction marker Claudin-5 (Supplements 1F-G). In addition, VeraVecs™ formed robust vessel-like structures when cultured in Matrigel (Supplement 1H), further validating their vessel formation capacity. VeraVecs™ were combined with dissociated organoid cells in a 1:1 ratio; notably, higher ratios of VeraVecs™ resulted in poor aggregate formation, while lower ratios resulted in insufficient VeraVec™ survival and lack of vascular structures (data not shown). In order to enhance the cellular complexity of our assembloid models, we also obtained hiPSC-derived CMPs, which were differentiated from hematopoietic progenitor cells as previously reported, and integrated them into our assembloid model via spontaneous migration (Figure 1K).^4^

**Figure 1.**
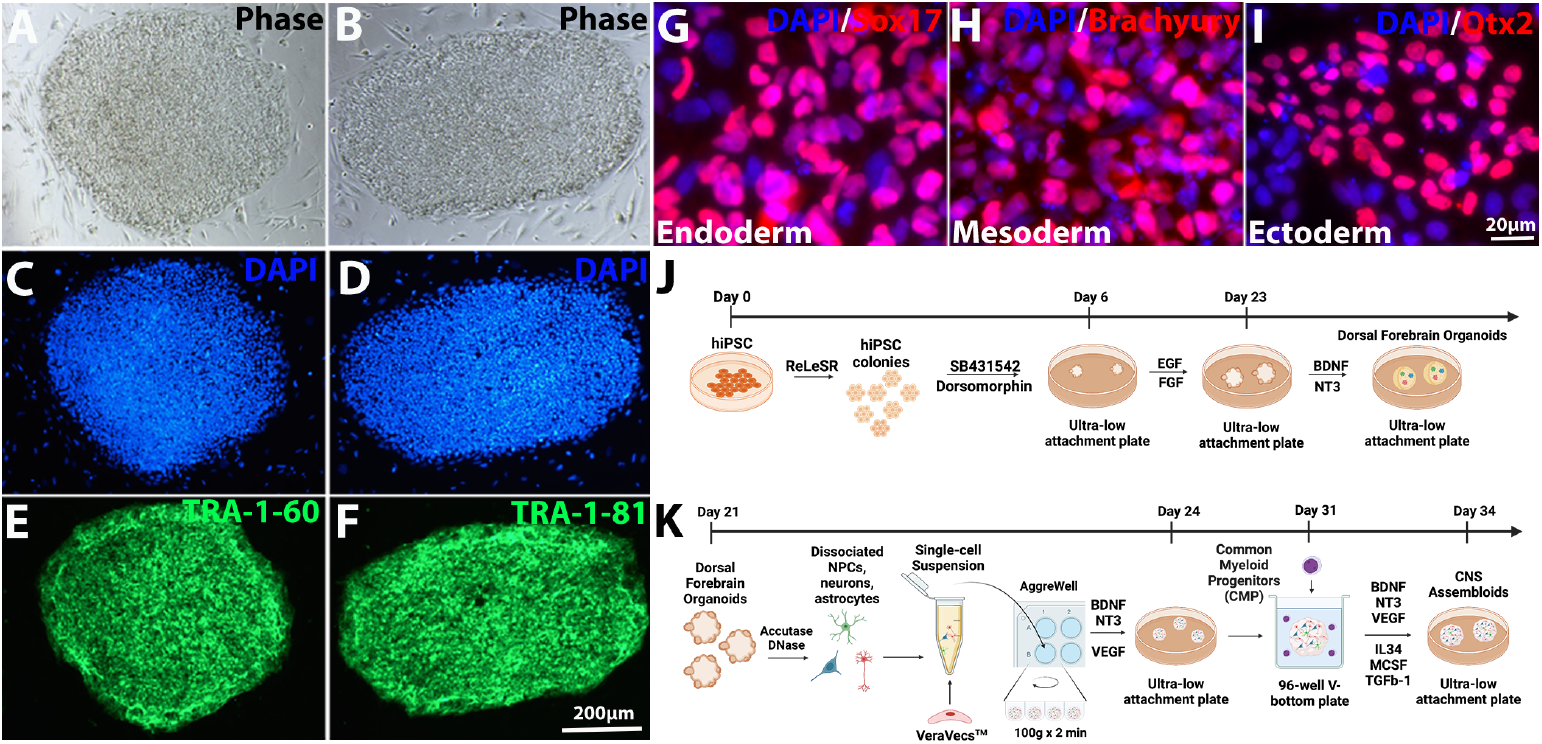
Characterization of human induced pluripotent stem cells (hiPSCs) and generation of dorsal forebrain organoids and vascularized assembloids. **(A-B):** Phase contrast image for hiPSC colonies. **(C-F):** Representative immunofluorescence images for **(C-D):** cell nucleus marker DAPI and pluripotency markers **(E):** TRA-1-60, and **(F):** TRA-1-81. **(G-I)** Representative immunofluorescence images for trilineage differentiation including **(G):** endoderm marker SOX17, **(H):** mesoderm marker Brachyury, and **(I):** ectoderm marker OTX2. **(J-K)** Schematics of the procedures used to generate **(J)** dorsal forebrain organoids and **(K):** vascularized assembloids.

Forebrain assembloids were characterized for populations of neuroectodermal cell types, displaying robust populations of NPCs with SOX2, neurons with NeuN, and astrocytes with GFAP, as seen in the organoid cultures (Figures 2A-B and Supplements 1A-D). We also demonstrated successful integration of VeraVecs™ into the assembloids and were able to detect small vessel-like structures in some areas of the assembloids (Figure 2D). Notably, the assembloids, which introduce the endothelial cell type into the 3D model, show a unique co-clustering pattern with concentrated astrocytes and endothelial cells (Figures 2C-E). This suggests that endothelial cells and astrocytes play critical roles in instructing each other’s subsequent development and organization and support other findings seen across multiple brain regions.^5-7^ Additionally, integrated CMPs were able to mature into microglia-like cells as verified by expression of microglia markers IBA1, TREM2, and CD11b, and showed robust process extension characteristic of surveilling microglia, validating this in-house method for facilitating immune-capable cell types (Figures 2F-H). In addition, we were also able to detect small numbers of oligodendrocytes, characterized by expression of oligodendrocyte marker MBP, in our assembloids (Figure 2I). To compare and evaluate the structural and organizational characteristics of our organoid and assembloid models, we conducted a comparison between 2.5-month-old assembloids and age-matched organoids for neuroepithelial loop distribution and size, astrocyte maturity, and synapse number. On average, the assembloids were found to have 3.02 more neuroepithelial loops per mm^2^ with 16.71% more proliferative area than organoids, suggesting that cells within 3-week-old organoids are not only robust enough to withstand dissociation but able to improve the coveted self-organizing character of 3D models with the addition of endothelial and microglial components (Figures 2J-L). GFAP expression in astrocyte processes was used to compare each model’s ability to promote astrocyte maturation. Given that GFAP is also moderately expressed in radial glia, we limited our analysis to GFAP positive cells located outside of the proliferative neuroepithelial regions and found that astrocyte processes contained within assembloids displayed a 79.59% higher GFAP intensity than those within organoids, further supporting the unique contribution of endothelial and microglial cells to astrocyte development (Figures 2M-O). We also found a significant increase (140.8 more synapses per 100 μm^2^) in the number of synapses in 2.5-month-old assembloids, compared with the age-matched organoids. Pre and post-synapsis were labeled by synaptophysin and PSD 95 antibodies, respectively (Figures 2P-Z). While organoids have been lauded for their ability to produce functional networks of neurons with utility in electrophysiological studies, assembloids may offer a more dynamic model, especially given the documented influence of non-neuronal cell types such as endothelial cells, astrocytes, and microglia on instructing synapse organization and network formation.^8,9^

**Figure 2.**
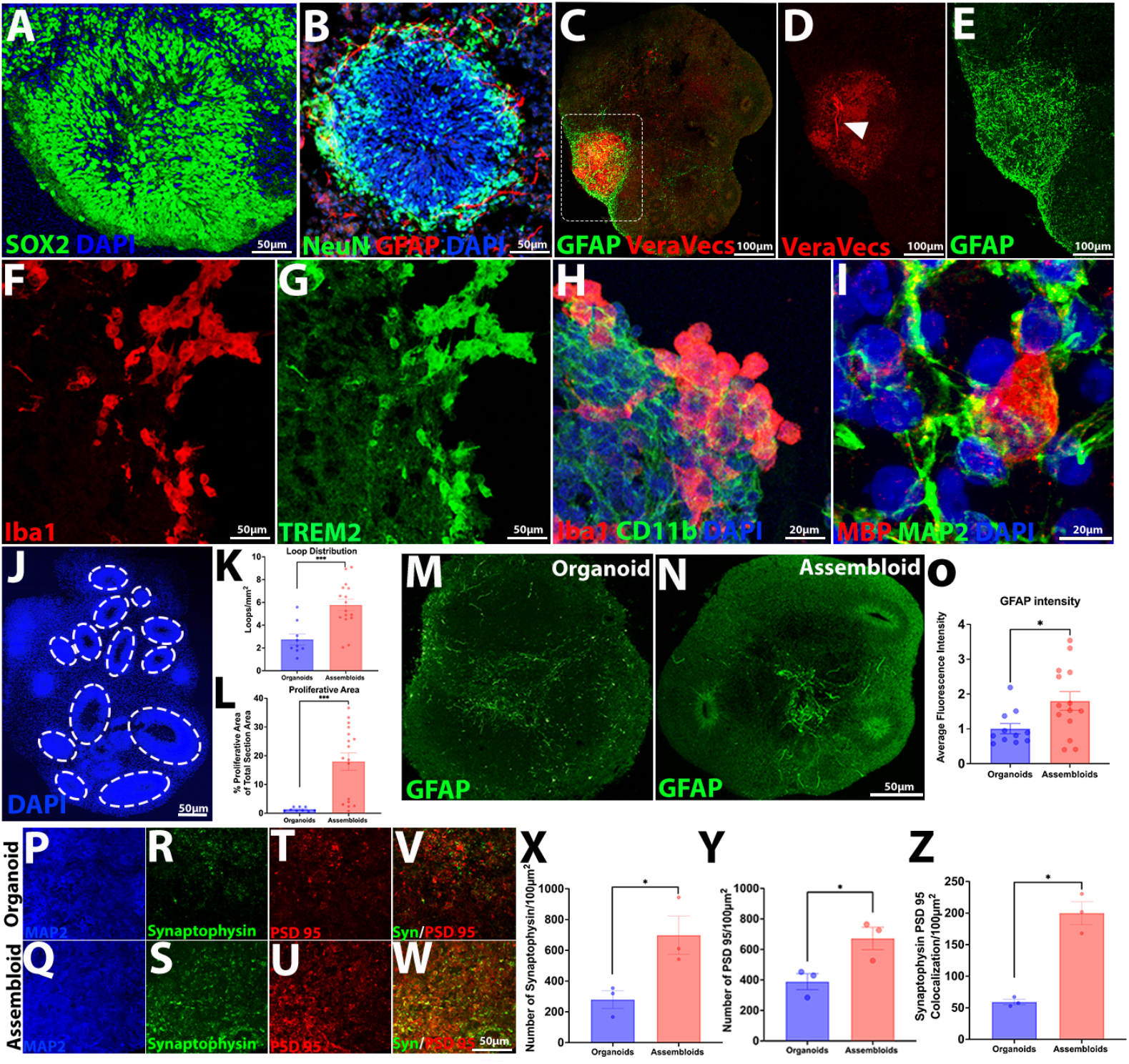
The vascularized assembloids contain multi-lineage cell types with unique organization with improved neuroepithelial loop features, astrocyte maturity, and synapse numbers. **(A-B):** Representative immunofluorescence images of neuroepithelial loop regions. **(A):** Neural progenitor cells (NPCs) marked with SOX2 make up most of the proliferative zone, along with surrounding cell nuclei marked with DAPI. **(B):** Neurons marked with NeuN and astrocytes marked with GFAP, along with cell nuclei marked with DAPI, emerge from proliferative zones, and migrate radially outward. **(C):** Representative immunofluorescence images of unique co-clustering between VeraVecs™ expressing red fluorescent protein (RFP) and astrocytes marked with GFAP. **(D-E):** Enlarged images show **(D)** emergence of vessel like structures, which are depicted with the arrowhead, and **(E)** robust GFAP expression in the same region. **(F-H):** Representative immunofluorescence images show presence of microglia-like cells expressing microglia markers **(F)** IBA1, **(G)** TREM2, and **(H)** CD11b. **(I):** Representative immunofluorescence image of oligodendrocytes marked with MBP contacting dendrites marked with MAP2. **(J):** Representative whole-section immunofluorescence image of neuroepithelial loop distribution in 2.5-month-old vascularized assembloids using DAPI. Outlines depict individual loops. **(K):** Bar graph showing mean loop distribution per mm^2^ ± SEM for organoids (n=9 sections from 9 organoids) and assembloids (n=16 sections from 7 assembloids). Assembloids had 3.02±1.01 more loops per mm^2^ (***p≤0.001). **(L):** Bar graph showing average percent proliferative area per section ± SEM for organoids (n=9 sections from 9 organoids) and assembloids (n=17 sections from 8 assembloids). Proliferative area is defined by area between lumen and outer region of each loop. Assembloids had 16.61±6.26% more proliferative area (***p≤0.001). **(M-N):** Representative whole-section immunofluorescence images of GFAP expression in 2.5-month-old **(M)** organoids and **(N)** assembloids. **(O):** Bar graph showing GFAP intensity in astrocyte processes ± SEM for organoids (n=11 processes) and assembloids (n=14 processes). Data is normalized to organoids. Assembloids showed 79.59±50.18% higher GFAP expression (*p≤0.05). **(P-W):** Representative immunofluorescence images of **(P-Q)** neural dendrite marker MAP2, **(R-S)** pre-synapse marker synaptophysin, **(T-U)** post-synapse marker PSD 95, and **(V-W)** synaptophysin and PSD 95 merge in 2.5-month-old **(P, R, T, V)** organoids and **(Q, S, U, W)** assembloids. **(X-Z):** Bar graphs showing **(X)** number of synaptophysin positive puncta, **(Y)** number of PSD 95 positive puncta, and **(Z)** colocalization of synaptophysin and PSD 95 per 100μm^2^ ± SEM for organoids (n=3 sections from 3 organoids) and assembloids (n=3 sections from 3 assembloids). Assembloids displayed significant increases in synaptophysin number by 418.71±107.59, PSD 95 number by 283.55±63.76, and the colocalization of synaptophysin and PSD 95 by 140.8±15.61 when compared to organoids (*p≤0.05).

To demonstrate the utility of this assembloid model in studying neurodegenerative diseases, we generated organoids and assembloids from hiPSCs harboring the tau^P301S^ mutation derived from a patient with Frontotemporal dementia and parkinsonism-linked to chromosome 17 (FTDP-17). Tau^P301S^ hiPSCs and their CRISPR-Cas9 corrected isogenic wild-type controls were karyotyped and sequenced to verify the presence of the point mutation and corrected base pair (Figures 3A-D). Wild-type and tau^P301S^ hiPSCs, along with non-isogenic wild-type VeraVecs™ and non-isogenic wild-type CMPs, were used to generate assembloids per the method illustrated in Figure 1J. We first measured the level of total and hyperphosphorylated tau proteins, identified using TauR1 and AT8 antibodies, respectively, in 2.5-month-old wild-type and tau^P301S^ assembloids to explore early differences in tau proteins. Compared with wild-type assembloids, tau^P301S^assembloids showed an enhanced total tau level by 86.16%, marked by TauR1 antibody, as well as an elevated phospho-tau level by 67.58%, marked by Tau AT8 antibody, suggesting an early manifestation in tau pathology (Figures 3E-L). To assess the effects of the tau^P301S^ mutation on astrocyte reactivity, we measured GFAP expression in astrocyte processes. We identified a 46.74% higher GFAP intensity in the processes of tau^P301S^ astrocytes than those of wild-type astrocytes (Figure 3M-N). These findings mirror previously reported findings of early astrocyte activation in PS 19 mice and suggest that astrocyte reactivity may be an early indicator of pathology.^10^ We also performed Sholl analysis on individual astrocytes to measure process extension and branching to determine whether we can also detect morphological changes in tau^P301S^ astrocytes. We didn’t detect any difference in longest process length or maximum branch intersections at any radius, suggesting that morphological changes are not present at this early time and perhaps do not occur as rapidly as changes in GFAP expression. (Figure 3M-P). Remarkably, elevated levels of GFAP were detected in 8-month-old tau^P301S^ organoids (data not shown), thereby indicating that this assembloid model could potentially reveal pathological phenotypes at an earlier stage. This significant finding holds great promise in facilitating the early identification of disease-relevant phenotypes, ultimately aiding in their timely detection.

**Figure 3.**
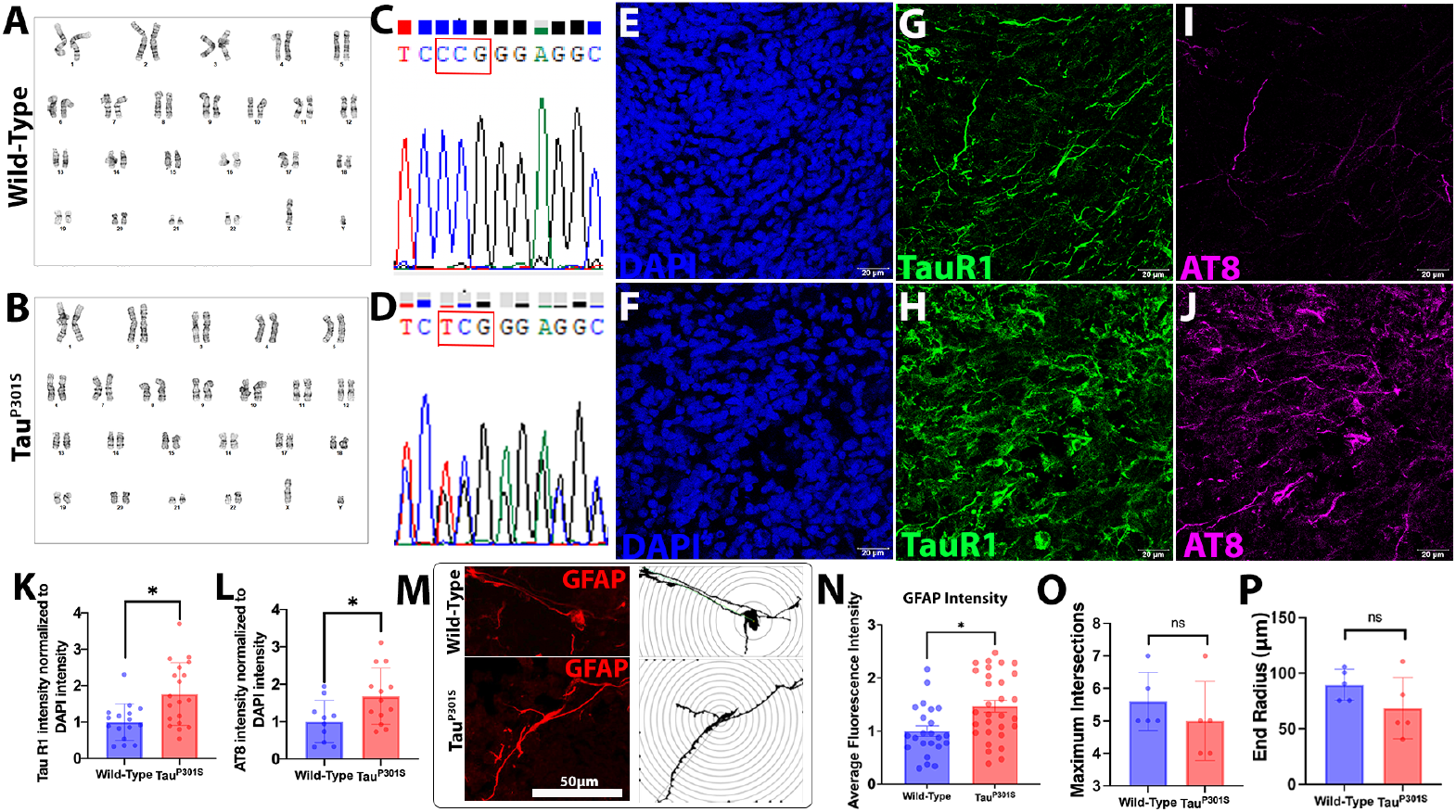
Tau^P301S^ assembloids display increased levels of total tau, hyperphosphorylated tau and GFAP. **(A-D):** hiPSCs harboring the tau^P301S^ and their isogenic wild-type controls were **(A-B)** karyotyped and **(C-D)** sequenced. **(E-J):** Representative immunofluorescence images of **(E-F)** DAPI, **(G-H)** total tau marked with TauR1, and **(I-J)** hyperphosphorylated tau marked with AT8 for 2.5-month-old assembloids generated from **(E, G, I)** wild-type and **(F, H, J)** tau^P301S^ hiPSCs. **(K):** Bar graph showing TauR1 intensity normalized to DAPI intensity ± SEM for wild-type (n=16) and tau^P301S^ assembloids (n=19). Data is normalized to the wild-type. Tau^P301S^ assembloids had 86.16±43.02% higher TauR1 expression (*p≤0.05). **(L):** Bar graph showing AT8 intensity normalized to DAPI intensity ± SEM for wild-type (n=10) and tau^P301S^ assembloids (n=13). Data is normalized to the wild-type. Tau^P301S^ assembloids had 67.58±36.17% higher AT8 expression (*p≤0.05). **(M):** Representative immunofluorescence images using GFAP and projections for individual astrocytes in 2.5-month-old wild-type and tau^P301S^ assembloids. Concentric radii are spaced 5μm apart. **(N):** Bar graph showing GFAP intensity in processes ± SEM for wild-type (n=23) and Tau^P301S^ astrocytes (n=30). Data is normalized to the wild-type. Tau^P301S^ astrocytes had 46.74±31.78% higher GFAP expression (*p≤0.05). **(O-P):** Bar graphs showing **(O)** maximum intersects at any radius and **(P)** end radius ± SEM for wild-type (n=5) and tau^P301S^ astrocytes (n=5). No significant difference was detected in either maximum intersects or end radius (ns: no significance).

The addition of microglia-like cells into the assembloids offers an improved model to probe neuroinflammation and study its role in neurodegenerative diseases. CMP-derived microglia-like cells incorporated within the tau^P301S^ assembloids showed no difference in overall size and area but displayed a more rod-like morphology when compared to those contained within wild-type assembloids, which were found to be more ramified (Figures 4A-E). Indeed, the rod-like morphology is an indication for early activated microglia and has been well documented in cases of neurodegeneration and injury.^11^ Microglia within tau^P301S^ assembloids were found to have a higher expression level of IBA1 by 62.93% than those in wild-type assembloids (Figure 4F). The observed augmentation of rod-like microglia and the upregulation of IBA1 expression in these microglia strongly indicate that a particular aspect of the mutated environment, such as elevated levels of total tau and hyperphosphorylated tau, is provoking an inflammatory response in otherwise healthy microglia. This suggests a potential link between the aberrant tau-related pathology and the pro-inflammatory activation of microglia in the studied context.^12^ To comprehensively evaluate the neuroinflammatory state of each model, we conducted the quantitative reverse transcriptase polymerase chain reaction (qRT-PCR), which allowed us to quantitatively assess the expression levels of key inflammatory markers, providing a robust and reliable measure of the neuroinflammatory condition in both models. Compared to wild-type assembloids, tau^P301S^ assembloids showed an elevated expression of the pro-inflammatory tumor necrosis factor-α (TNF-α), while no difference was detected in another pro-inflammatory factor, complementary factor B (CFB) (Figure 4G-H). Furthermore, the tau^P301S^ assembloids also showed a diminished expression of the anti-inflammatory marker S100A10 (Figure 4I). Together, this morphological and transcriptomic data suggest that the tau^P301S^ mutation, whether through neurons or astrocytes, or both, pushes microglia into an increasingly activated state, even at a relatively early timepoint and is in line with other findings in the field of tauopathy, which suggest that microglia may exist in a chronically inflamed state in disease conditions.^13^ It is noteworthy that we did not observe significant expression of inflammatory markers in either wild type or tau^P301S^ organoids, likely due to their lack of microglia-like cells, in turn validating the immune competency and capability of this improved assembloid system (Figure 4G-I).

**Figure 4.**
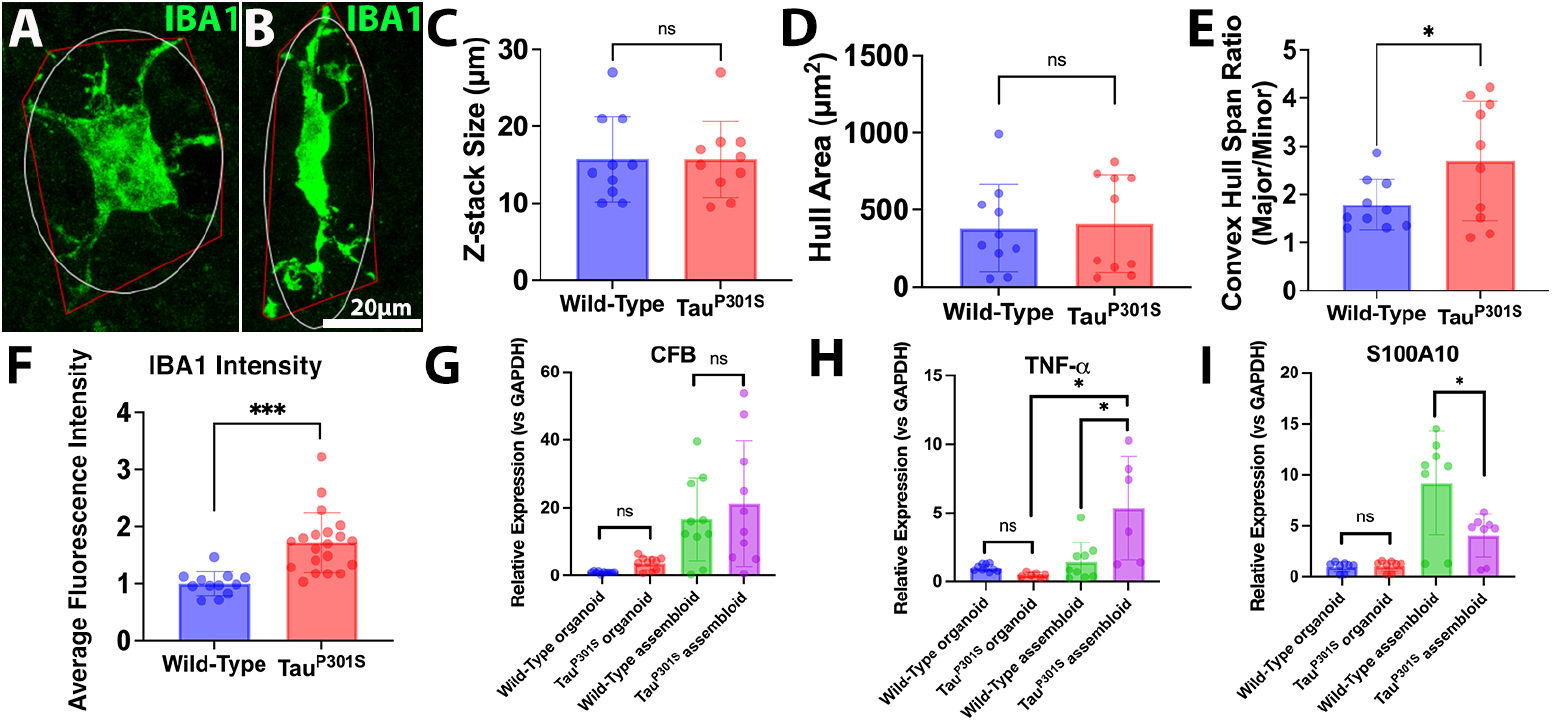
Tau^P301S^ assembloids induce enhanced neuroinflammation and microglial activation. **(A-B):** Representative immunofluorescence images for **(A)** ramified and **(B)** rod-like microglia marked with IBA1 in 2.5-month-old wild-type and tau^P301S^ assembloids. Red outlines represent convex hull area and white outlines represent fitted ellipse used to calculate convex hull span ratio. **(C-D):** Bar graphs showing **(C)** z-stack size and **(D)** hull area ± SEM for microglia in wild-type (n=10) and tau^P301S^ assembloids (n=10). No significant difference was detected in either z-stack size or hull area (ns: no significance). **(E):** Bar graph showing convex hull span ratio ± SEM for microglia in wild-type (n=10) and tau^P301S^ assembloids (n=10). Convex hull span ratio is calculated by taking the ratio of the major and minor axis of the ellipse fitted to the convex hull area. Microglia in tau^P301S^ assembloids had a 0.90±0.62 larger ratio and thus a more rod-like morphology (*p≤0.05). **(F):** Bar graph showing IBA1 intensity ± SEM for microglia in wild-type (n=12) and Tau^P301S^ assembloids (n=20). Data is normalized to the wild-type. Microglia in tau^P301S^ assembloids had a 62.93±26.6%) higher expression of IBA1 (***p≤0.001). **(G-I):** Bar graphs depicting relative expression of **(G)** CFB, **(H)** TNF-α, and **(I)** S100A10 to GAPDH in 2.5-month-old wild-type (n=10; n=10; n=8) and tau^P301S^ organoids (n=10; n=6; n=8) and wild-type (n=10; n=9; n=8) and tau^P301S^ assembloids (n=10; n=6; n=8). No significant difference was detected between assembloids in CFB expression (ns: no significance). Tau^P301S^ assembloids had a 3.09±1.89 fold increased expression of TNF-α and a 5.14±1.06 fold decreased expression of S100A10 compared to wild-type assembloids (*p≤0.05 using student’s t-test). Tau^P301S^ assembloids had a 3.67±1.89 increased expression of TNF-α compared to tau^P301S^ organoids (*p≤0.05 using ANOVA).

## Discussion and Conclusion

Our study successfully developed a sophisticated human CNS model that integrates crucial cellular elements often absent in current CNS organoid production, thereby enhancing its relevance to CNS development and diseases. Indeed, the incorporation of microglia-like cells and endothelial cells in the assembloids not only resulted in enhanced cell growth and increased maturity but also more accurately recapitulated the phenotypic changes observed in tauopathies. Interestingly, tau^P301S^ assembloids exhibited notable increases in pro-inflammatory phenotypes among astrocytes and microglia, which were not fully evident in the organoids, supporting findings that suggest the critical role of glia in advancing the disease.^14,15^ The inclusion of microglia-like cells derived from a separate wild-type hiPSC line in our assembloid models revealed a non-autonomous microglial response to the tau pathology observed in neurons, suggesting potential involvement of astrocytes and the surrounding microenvironment in driving this response. We are keenly interested in further advancing our research by developing assembloids derived from a single hiPSC line, allowing for more accurate disease modeling that closely represents the clinical context of patients. While our assembloid models have achieved successful integration of vascular endothelial cells and microglia-like cells, there is a need for further advancements to optimize the formation of vascular structures within these models and to augment the robust presence of myelinating oligodendrocytes, which is currently constrained in our study. Strikingly, in various tauopathy models, the presence of pathological tau species has been detected not only in neurons but also in astrocytes, microglia, and vascular endothelial cells, indicating the potential role of these cell types as mediators in the propagation of pathological tau.^16,17^ However, an alternative perspective suggests that these cells may indeed engage in a protective response, actively involved in the breakdown and clearance of pathological tau, as previously proposed.^18,19^ In summary, our human assembloid system represents a significant advancement in modeling neurological disorders, providing a valuable tool to investigate the intricate intercellular mechanisms and to gain deeper insights into these complex processes.

## Acknowledgements

We thank Dr. Celeste Karch for providing us with the tau^P301S^ hiPSCs and the CRISPR-Cas9 edited wildtype hiPSCs. We thank Angiocrine Inc. and Dr. Daniel Nolan for providing us with VeraVecs™, as well as their insight in microvasculature. We thank Dr. Nicholas Kanaan for providing us with TauR1 antibodies. We also thank Yash Agarwal for help with the qRT-PCR experiments.

## Data Availability

The raw data supporting the conclusions of this article will be made available by the authors, without undue reservation.

