## Supplementary Materials for "Vascularized Brain Assembloids with Enhanced Cellular Complexity Provide Insights into The Cellular Deficits of Tauopathy"

### Methods and Materials

#### *hiPSC Culture*

Patient-derived hiPSCs harboring the tau$^{P301S}$ mutation and their CRISPR-Cas9 corrected isogeneic controls were obtained from the Karch Lab at Washington University School of Medicine. hiPSC's were cultured in 5% CO2 at 37 °C in mTeSR Plus (STEMCELL Technologies, 100-0276) as previously reported.[1] hiPSCs and their derivatives were fixed with 4% paraformaldehyde (PFA) and further validated for their pluripotency and three germ-layer differentiation by immunocytochemistry (ICC) using Pluripotent Stem Cell 4-Marker ICC Kit (Invitrogen) and Human Three Germ Layer 3-Color ICC Kit (R&D Systems).

#### *VeraVec™ Culture*

Cryopreserved RFP-labelled VeraVecs™ were obtained from Angiocrine Inc. VeraVecs™ were cultured in serum-free conditions and supplemented with VEGF (50 ng/mL; PeproTech, 100-20) and bFGF (20 ng/mL; PeproTech, 100-18B) as previously reported.[2] Full medium changes were performed every other day. Upon >90% confluency, cells were lifted for replating, freezing, or assembloid generation using accutase (STEMCELL Technologies, 07920). VeraVecs™ were fixed with 4% PFA and immunostained with anti-CD31 (Thermo Fisher, 11-0319-41) and anti-Claudin-5 (BiCell, 00205).

#### *Dorsal Forebrain Cortical Organoid Culture*

Dorsal forebrain organoids were generated from tau$^{P301S}$ and isogenic control hiPSC lines per a modified version of Pasca's protocol.[3] Upon reaching ~75% confluency, hiPSCs were lifted using ReLeSR (STEMCELL Technologies, 100-0483), and 1 well of hiPSCs was plated into 1 well of a 6-well ultra-low attachment plate in mTeSR Plus with ROCK inhibitor Y-27632 (10μM; Tocris, 1254), Dorsomorphin (5 μM; Sigma, C956T21), and SB-431542 (10μM; Tocris, 1614) to promote embryoid bodies (EBs) formation and neuronal induction. A full medium change without ROCK inhibitor Y-27632 was performed on Day 3. From Day 6 until dissociation, EBs were placed onto an orbital shaker (60 rpm) and medium was changed to neuronal differentiation medium (NM) prepared using Neurobasal A (Thermo Fisher, 10-888-022), B-27 without Vitamin A (Thermo Fisher, 12587010), and Glutamax (Thermo Fisher, 35-050-061). NM was supplemented with EGF (20ng/mL; PeproTech, AF-100-15) and bFGF (20ng/mL). On Day 23, medium was switched to NM supplemented with BDNF (20 ng/mL; PeproTech, 450-02) and NT3 (20 ng/mL; PeproTech, 450-03). On Day 45, NM was also supplemented with B-27 with Vitamin A (Thermo Fisher, A3582801). All organoids were cultured in 5% $CO_2$ at 37 °C on an orbital shaker. Medium changes were performed every 3-4 days.

*Assembloid Generation*

1. Organoid Dissociation

Day 14-21 organoids were collected and transferred into 15 mL falcon tubes. After letting organoids sink and removing supernatant, 5 mL accutase and 150 μL DNase (Sigma, DN25) were added and mixed well with samples. Tubes were placed in a 37 °C water bath for 20 minutes, with gentle tapping to mix solution every 5 minutes. Tubes were then transferred to room temperature, samples were allowed to sink, and supernatant was removed. 5x3-minute washes using Neurobasal A medium were performed, ensuring to resuspend loosely attached organoids during each wash to remove any residual enzyme solution. After the fifth wash, cells were resuspended in NM medium.

A slightly harsh trituration using a P1000 tip was performed to create a single-cell suspension and cells were subsequently counted using a hemocytometer.

2. AggreWell Preparation

AggreWell plates were prepared in parallel to cell preparation steps per the product information sheet made available by STEMCELL Technologies. Anti-adherence solution (STEMCELL Technologies, 07010) was added to each well of a 24-well AggreWell plate and centrifuged in a swinging rotor-bucket at 1300 x g for 5 minutes. After ensuring no bubbles remained trapped in microwells, solution was aspirated, and wells were washed with warm Neurobasal A.

3. Incorporation of VeraVecs™ into the Assembloids

Aggregate solutions were made in 1.5mL Eppendorf tubes by combining organoid-dissociated cells and VeraVecs™ in a 1:1 ratio to reach a density of 1.5 million cells per well. Tubes were topped up to 1mL with NM medium, and following gentle trituration with P1000, cell solutions were plated dropwise in an additional 1mL of medium to achieve a seeding volume of 2mL per well. NM medium was supplemented with ROCK inhibitor Y-27632 (10μM), BDNF (20 ng/mL), NT3 (20 ng/mL), and VEGF (10 ng/mL). AggreWell plates were immediately centrifuged at 100 x g for 2 minutes. Samples were then observed under the microscope to ensure even and proper seeding in microwells, and plates were then carefully placed into incubator and left undisturbed for 72 hours to allow seeded cells to form robust aggregates. After 72 hours, aggregates were transferred to low attachment plates and full medium change was performed excluding ROCK inhibitor Y-27632. Aggregates were allowed to recover for 1 week before addition of common myeloid progenitors (CMPs).

4. Addition of Common Myeloid Progenitors to Form Assembloids

Wild type hiPSCs were matured into hematopoietic progenitor cells (HPCs) and sorted into CD45+/CD18+ CMPs as previously reported.[4] Cryopreserved CMPs were obtained from the Human Stem Cell Core at the Children's Hospital of Philadelphia. CMPs were thawed and resuspended in NM medium supplemented with BDNF (20 ng/mL), NT3 (20 ng/mL), VEGF (10 ng/mL), IL-34 (100 ng/mL; PeproTech, 200-34), TGFβ-1 (50 ng/mL; PeproTech, 100-21), and M-CSF (25 ng/mL; PeproTech, 300-25). CMPs were then added to 96-well V-bottom plates at a density of 25,000 cells per well in a seeding volume of 150uL and combined with one aggregate (see previous section), and the resulting co-culture was incubated for an additional 72 hours. After incubation period, "assembloids" were transferred to 6-well low-attachment plates and placed on an orbital shaker (60 rpm) for long term culture. Medium changes were performed every 3-4 days.

*Immunofluorescence Analyses*

Samples were fixed in 4% PFA for 16 hours at 4°C and saturated with 30% sucrose solution before embedding in M1 embedding matrix (Thermo Fisher, 1310). Frozen samples were sectioned in 35µm slices and stained as previously reported.[5] Primary antibodies were sourced as follows: SOX2 (Sigma, AB5603), NeuN (Abcam, ab104224), GFAP (Abcam, ab53554), IBA1 (ABclonal, A19776), TREM2 (R&D Systems, AF1828), CD11b (Proteintech, 66519-1-Ig), MBP (ABclonal, A1664), AT8 (Thermo Fisher, MN1020), Synaptophysin (Proteintech, 60191-1-AP), PSD 95 (Cedarlane labs, 124008(SY)), and MAP2 (Novus, NB300-213). Anti-TauR1 was obtained from the Kanaan Lab at Michigan State University. Appropriate goat or donkey secondary antibodies conjugated to Alexa Fluor 405, 488, 555, and 647 were used. All images were acquired using either

20x oil or 63x oil objectives on a Leica SP8 Confocal Microscope or a Zeiss AxioObserver Z1 Microscope. Image processing was done using Fiji and Photoshop2023.

*RNA Isolation and qRT-PCR Analyses*

RNA was extracted from samples using the Ambion PureLink RNA Mini Kit (Thermo Fisher) and quality was assessed using the NanoDrop spectrophotometer. Reverse transcription was done using the High-Capacity cDNA Reverse Transcription Kit (Applied Biosystems) to synthesize cDNA. qRT-PCR was performed on the StepOnePlus™ Real-Time PCR System (Applied Biosystems) using the 2X Universal SYBR Green Fast qPCR Mix (ABclonal). RNA primers were obtained from IDT DNA with sequences listed below:

CFB Forward: 5’ CTC TTG TCT GGA GGT GTG ACC 3’

CFB Reverse: 5’ CCG CCT TTG ATC TCT ACC CC 3’

TNF-α Forward: 5’ AGC CCA TGT TGT AGC AAA CCC 3’

TNF-α Reverse: 5’ GGA CCT GGG AGT AGA TGA GGT 3’

S100A10 Forward: 5’ TTC GCT GGG GAT AAA GGC TAC 3’

S100A10 Reverse: 5’ CAG AGG GTC TTT TTG ATT TTC CA 3’

*Statistical Analyses*

Data was initially recorded in an Excel sheet, and GraphPad Prism9 was subsequently employed to generate graphs. The outliers in the qRT-PCR data were removed using GraphPad Prism9. Values are reported as means ± SEM. A students t-test was used for comparison of two groups while a one-way ANOVA was used for comparison of more than two groups.

1. Yates PL, Patil A, Sun X, et al. A cellular approach to understanding and treating Gulf War Illness. *Cell Mol Life Sci*. Nov 2021;78(21-22):6941-6961. doi:10.1007/s00018-021-03942-3

2. Seandel M, Butler JM, Kobayashi H, et al. Generation of a functional and durable vascular niche by the adenoviral E4ORF1 gene. *Proceedings of the National Academy of Sciences*. 2008/12/09 2008;105(49):19288-19293. doi:10.1073/pnas.0805980105

3. Sloan SA, Andersen J, Pașca AM, Birey F, Pașca SP. Generation and assembly of human brain region-specific three-dimensional cultures. *Nat Protoc*. 2018;13(9):2062-2085. doi:10.1038/s41596-018-0032-7

4. Mills JA, Paluru P, Weiss MJ, Gadue P, French DL. Hematopoietic differentiation of pluripotent stem cells in culture. *Methods Mol Biol*. 2014;1185:181-94. doi:10.1007/978-1-4939-1133-2_12

5. Yates PL, Case K, Sun X, Sullivan K, Baas PW, Qiang L. Veteran-derived cerebral organoids display multifaceted pathological defects in studies on Gulf War Illness. Original Research. *Frontiers in Cellular Neuroscience*. 2022-December-23 2022;16doi:10.3389/fncel.2022.979652

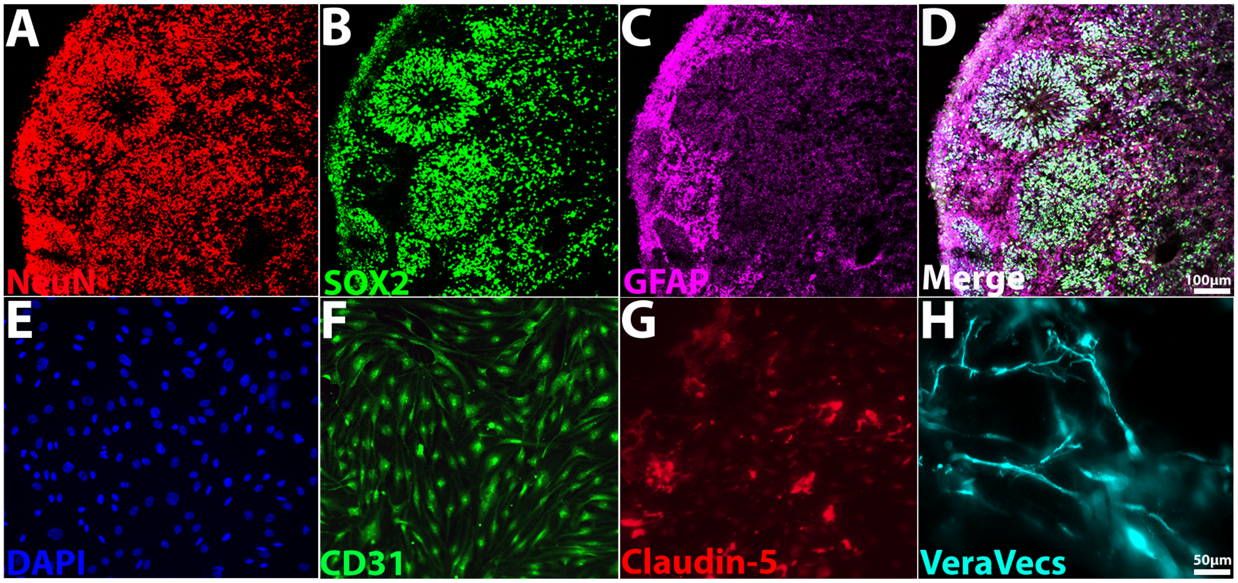


**Supplementary Figure 1. Characterization of dorsal forebrain organoids and VeraVecs™.** **(A-D):** Representative immunofluorescence images of neuroepithelial loop regions and cellular organization including **(A)** neurons marked with NeuN, **(B)** NPCs marked with SOX2, and **(C)** astrocytes marked with GFAP. **(D):** Neurons and astrocytes emerge from these proliferative loop regions and take up residence in surrounding cortical regions. **(E-G):** Representative immunofluorescence images of VeraVecs™ including **(E)** DAPI, **(F)** endothelial cell marker CD31, and **(G)** tight junction marker Claudin-5. **(H):** Representative multi-plane immunofluorescence image of VeraVecs™ expressing RFP organizing into vessel-like structures when cultured in Matrigel.
